# Galactan mobilization during carbon starvation compromises cell wall-mediated fungal resistance in Arabidopsis

**DOI:** 10.1101/2025.05.28.656597

**Authors:** Louisa Reckleben, Richard N. Morton, Kristina S. Munzert-Eberlein, Sabine Thiel, Carsten Rautengarten, Viviana Tan, Klara L. Harres, Lena Zanoni, Berit Ebert, Timo Engelsdorf

**Affiliations:** Molecular Plant Physiology, Department of Biology, Philipps-Universität Marburg, 35043 Marburg, Germany; Faculty of Biology and Biotechnology, Ruhr University Bochum, 44801 Bochum, Germany

**Keywords:** Cell wall, pectin, galactose, sugar salvage, starch, phytopathogenic fungi, pathogen resistance, pathogen defense, *Arabidopsis thaliana*, *Colletotrichum higginsianum*

## Abstract

Polysaccharides are the main components present in plant cell walls. They form a network that is dynamically modified during growth and upon both abiotic and biotic stress. We investigated how the cell wall of Arabidopsis rosettes is remodeled during periods of dark-induced starvation in the wild type and in *plastidic phosphoglucomutase* (*pgm*) mutants, which suffer from periodic starvation due to starch deficiency. Time-course analysis demonstrated that up to one fifth of the galactose present in leaf cell walls is reversibly released upon starvation, while other cell wall monosaccharides were less affected. An investigation of *BETA-GALACTOSIDASE* (*BGAL*) expression and the analysis of *bgal* mutants indicated that *BGAL1* and *BGAL4* contribute to the release of cell wall galactose upon starvation. Increased transcript abundance of *UDP-GLUCOSE 4-EPIMERASE* (*UGE*) *1* and *3* under starvation proposed an increased flux through the galactose salvage pathway, however an analysis of the UDP-galactose pool in mutant plants indicated redundancy with other UGEs. Simultaneously to galactan degradation, *GALACTAN SYNTHASE1* (*GALS1*) expression was reduced, attenuating the synthesis of new galactan chains. We show that overexpression of GALS1 prevents depletion of the recyclable cell wall galactose pool and is sufficient to rescue impaired penetration resistance to the hemibiotrophic fungal pathogen *Colletotrichum higginsianum* upon dark-induced and periodic starvation. Our data suggest that pectic galactan in the plant cell wall serves as a sugar resource during starvation conditions. However, galactose release from the wall leads to impaired penetration resistance against a fungal pathogen, causing a tradeoff between sugar supply for plant metabolism and preformed defense.

**Significance statement:** Polysaccharides present in plant cell walls must be dynamically modified in response to external stimuli, but it is not well understood how cell wall remodeling is adjusted in response to metabolic cues and how this affects defense against pathogens. Here, we report that galactose is reversibly released from cell walls upon carbon starvation, which critically affects penetration resistance to a fungal pathogen.

## Introduction

Cell walls provide support to plant cells and delimit them against neighboring cells and their environment. Polysaccharides represent the main components of the wall and form a network of cellulose, hemicelluloses and pectins. This network is dynamically remodeled during growth and development, as well as in response to stress (Delmer *et al*., 2024). When carbon supply from photosynthesis is limited, alterations in cell wall composition occur that might be linked to carbon provision from cell wall polymers. Dark-induced sugar starvation in detached *Arabidopsis thaliana* (hereafter: Arabidopsis) leaves leads to the induction of glycoside hydrolases and to reduced amounts of pectin and hemicellulose after two days of starvation (Lee *et al*., 2007). Carbon depletion of intact Arabidopsis plants caused by an extended dark phase leads to distinct transcriptional changes aimed at adapting metabolism to the low carbon availability (Usadel *et al*., 2008). Responses are similar in starch-deficient plastidic *phosphoglucomutase* (*pgm*) mutants at the end of the night, when they suffer carbon limitation after soluble sugars are depleted (Caspar *et al*., 1985, Gibon *et al*., 2004, Usadel *et al*., 2008, Engelsdorf *et al*., 2013). This periodic nocturnal carbon limitation in *pgm* causes pronounced alterations in leaf cell wall composition. Cell wall glycan profiling with monoclonal antibodies indicated a reduced presence of epitopes from pectic rhamnogalacturonan-I (RG-I) side chains and cell wall monosaccharide analysis revealed reduced relative amounts of arabinose and galactose in the *pgm* mutant (Engelsdorf *et al*., 2017).

RG-I consists of a backbone of alternating galacturonic acid and rhamnose units decorated with arabinan, galactan and arabinogalactan side chains and can constitute up to one third of all pectins in leaf primary cell walls (Mohnen, 2008). The remaining pectin is comprised of homogalacturonan (up to two thirds), consisting entirely of galacturonic acid, and of substituted homogalacturonan such as RG-II and xylogalacturonan (Delmer *et al*., 2024). Nucleotide sugars, primarily made in the cytosol, serve as substrates for polysaccharide formation (Bar-Peled and O’Neill, 2011). UDP-D-glucose represents the central intermediate in the nucleotide sugar interconversion pathway and can be directly interconverted to UDP-D-glucuronic acid, UDP-L-rhamnose and UDP-D-galactose (Bar-Peled and O’Neill, 2011). In Arabidopsis, interconversion of UDP-D-glucose and UDP-D-galactose is catalyzed by a family of five UDP-glucose 4-epimerases (UGEs), with the expression of *UGE1* and *UGE3* being co-regulated with carbohydrate catabolism and that of *UGE2*, *UGE4* and *UGE5* with carbohydrate biosynthesis (Barber *et al*., 2006). UDP-rhamnose/UDP-galactose transporters (URGTs) translocate the substrates for galactan biosynthesis into the Golgi lumen (Rautengarten *et al*., 2014). After import into the Golgi, nucleotide sugars are used as substrates by glycosyltransferases to assemble cell wall polysaccharides. RG-I:Rhamnosyltransferase (RRT) and RG-I:Galacturonosyltransferase (RGGAT) build the RG-I backbone, while UDP-D-galactose serves as substrate for the synthesis of β-1, 4-galactan catalyzed by the galactosyltransferases GALACTAN SYNTHASE 1 (GALS1), GALS2 and GALS3 (Liwanag *et al*., 2012, Ebert *et al*., 2018, Takenaka *et al*., 2018, Amos *et al*., 2022). GALS1 is furthermore able to regulate the length of galactan chains by terminally adding arabino*pyranose* (Ara*p*) (Laursen *et al*., 2018). The nucleotide sugar UDP-L-Ara*p* is formed in the Golgi via sequential conversion of UDP-D-glucose to UDP-D-glucuronic acid and UDP-D-xylose and the subsequent activity of UDP-xylose 4-epimerases (UXEs) (Mariette *et al*., 2021). In Arabidopsis, MURUS4 (MUR4) represents the main Golgi-localized UXE isoform, whereas UGE1 and UGE3 exhibit additional cytosolic UXE activity (Burget *et al*., 2003, Kotake *et al*., 2009, Umezawa *et al*., 2024). In the cytosol, UDP-L-Ara*p* is converted to UDP-L- arabino*furanose* (UDP-L-Ara*f*) by mutases before being moved back into the Golgi by UDP-Ara*f* transporters (UAfTs) where Ara*f* serves as the predominant building block for synthesis of RG-I α-1, 5- arabinan, putatively via ARABINAN DEFICIENT 1 (ARAD1) (Harholt *et al*., 2006, Rautengarten *et al*., 2011, Rautengarten *et al*., 2017). After assembly in the Golgi, cell wall polymers like pectins are secreted into the apoplast and integrated into the cell wall (Hoffmann *et al*., 2021). Nucleotide sugars can also be formed from recycled cell wall sugars that are released from cell wall polysaccharides and transported into the cytosol by sugar transport proteins (Barnes and Anderson, 2018). Metabolic salvage of these sugars is achieved via the activities of sugar-1-kinases and UDP-sugar-pyrophosphorylase (USP) (Geserick and Tenhaken, 2013). Galactose salvage is initiated by phosphorylation through galactokinase (GALK) (Egert *et al*., 2012). Subsequently, galactose-1-phosphate is converted to UDP-D-galactose by USP, while conversion to UDP-D-glucose seems to be mainly achieved by UGE1 and UGE3 (Barber *et al*., 2006, Geserick and Tenhaken, 2013).

As the outer most layer of plant cells, cell walls represent an important barrier against phytopathogen infection and changes in cell wall composition can affect pathogen resistance (Molina *et al*., 2021). Cell wall-penetrating fungal pathogens are adapted to the cell wall composition of their hosts by expressing cell wall-degrading enzymes that allow local cell wall decomposition and fungal entry (for a review see Munzert and Engelsdorf, 2025). The hemibiotrophic ascomycete *Colletotrichum higginsianum* infects Arabidopsis rosettes by penetrating the leaf surface with a specialized infection structure termed appressorium. Appressoria use a combination of turgor pressure and cell wall-degrading enzyme activity to drive a penetration peg through the epidermal cuticle and cell wall (Ryder *et al*., 2022). After successful entry around 36 hours post infection, *C. higginsianum* develops biotrophic hyphae in the leaves, but remains separated from the host cytoplasm by the host plasma membrane (O’Connell *et al*., 2004). Around 3 days post infection, the fungus switches to a necrotrophic lifestyle, leading to lysis of infected cells and colonization of adjacent leaf tissue (Engelsdorf *et al*., 2013). The entry rate of *C. higginsianum* strongly depends on pectin content and/or composition of the host cell walls, likely explaining enhanced fungal entry into *pgm* leaves (Engelsdorf *et al*., 2017).

Here, we investigated how carbon starvation leads to dynamic changes of the cell wall during extended nights in wild type and *pgm* rosettes and assessed the importance of cell wall remodeling for resistance to fungal infection. We found that galactose is reversibly released from cell wall galactan upon carbon starvation and that the abundance of cell wall galactan determines the entry rate of *C. higginsianum*.

## Results

### Starch-deficiency and dark-induced starvation cause similar rosette cell wall phenotypes

Our previous results suggested that the altered cell wall composition and the reduced abundance of RG-I side chain epitopes in *pgm* rosettes are caused by periodic nocturnal carbohydrate starvation in diurnal light / dark cycles (Engelsdorf *et al*., 2017). To test if dark-induced starvation in the wild type can phenocopy the cell wall composition of *pgm*, we subjected wild type (Col-0) and *pgm* plants to an extended night (XN) and sampled rosettes for analyses of the cell wall monosaccharide composition after 12, 24, 48 and 72 hours from the end of the regular night (EN) of a 12-hour light / 12-hour dark cycle. Starch reserves in the wild type were depleted after 4 hours of XN and soluble sugar content of both genotypes reached a minimum after 12 to 24 hours of XN (Fig. S1). In Col-0, we observed a significantly reduced cell wall galactose content after 12 hours XN (Fig. 1a). Galactose content was further reduced with prolonged darkness and, at 24 to 48 hours XN, it resembled the galactose content in *pgm* rosette cell walls at EN (Fig. 1a,b). After 72 hours XN, galactose levels in wild type cell walls were reduced by 18% compared to those before onset of XN (32.6 mol% at EN vs 26.6 mol% at XN 72h; Fig. 1a). While arabinose was also identified as being less abundant in *pgm* cell walls (Engelsdorf et al., 2017), in Col-0 it was only reduced after 72 hours XN and remained more abundant than in *pgm* cell walls (Fig. 1a,b). Cell walls of *pgm* rosettes contained less galactose after 24 hours of XN but exhibited no further decline until 72 hours of XN (Fig. 1b). Analysis of *gals1/2/3* mutant cell walls showed 23% less galactose compared to Col-0 and no further reduction upon XN treatment, indicating that only the GALS-dependent galactose pool is released under dark-induced starvation (Fig. S2). Reduced amounts of galactose were accompanied by a relative increase of xylose (Col-0) and rhamnose (Col-0 and *pgm*; Fig. 1a,b). Overall, the cell wall monosaccharide composition in Col-0 showed the highest degree of similarity to *pgm* EN cell walls after 72 hours XN (Fig. 1a,b). To test if the starvation-dependent reduction in cell wall galactose content can be reverted after XN, we subjected Col-0 plants to 72 hours XN before returning them to a regular light/dark cycle. While galactose content was reduced to 83% of that of the EN after XN, it recovered to 96% at 72 hours after return to a regular light/dark cycle (Fig. 1c).

**Figure 1.**
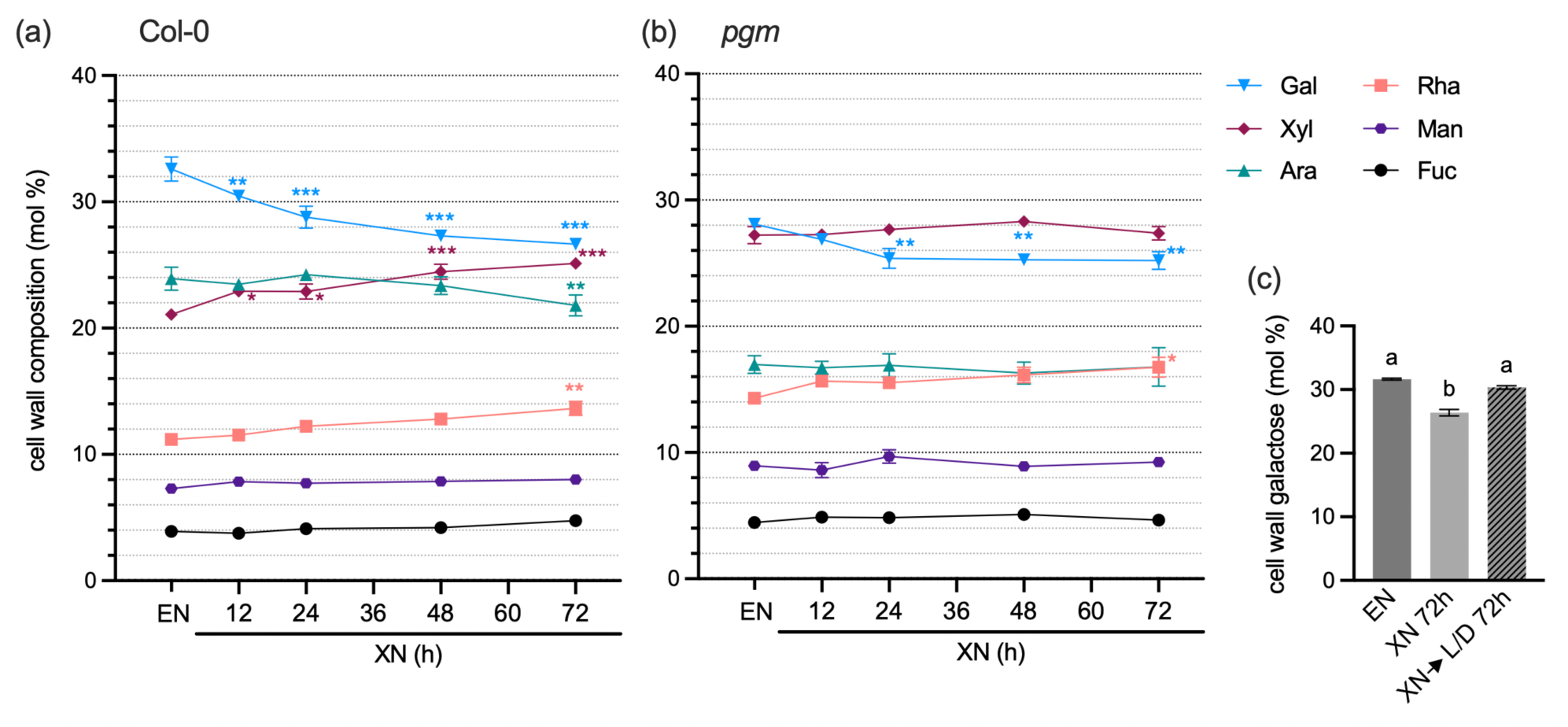
: Galactose amount in leaf rosette cell walls is reversibly decreased upon dark-induced starvation. (a) Col-0 and (b) *pgm* plants were subjected to an extended night (XN) after a diurnal 12h light / 12h dark phase. Whole rosettes were sampled at the end of the regular night (EN) and after 12, 24, 36, 48, 60 and 72h XN. The relative composition of the neutral cell wall monosaccharides galactose (Gal), xylose (Xyl), arabinose (Ara), rhamnose (Rha), mannose (Man) and fucose (Fuc) is depicted as the molar percentage of total cell wall neutral monosaccharide content (mol %). Values are means (n=4) and error bars represent the SEM. Asterisks indicate statistically significant diXerences to EN samples for each monosaccharide according to two-way ANOVA and Dunnett’s multiple comparisons test (*p<0.05, **p<0.01, ***p<0.001). (c) Molar percentage of galactose in cell wall neutral monosaccharides at EN, after 72h XN and after 72h XN followed by 72h in a 12h/12h light/dark (L/D) cycle. Values are means (n=4) and error bars represent the SEM. DiXerent letters indicate statistically significant diXerences according to one-way ANOVA and Tukey’s multiple comparisons test (α = 0.05).

Taken together, dark-induced starvation in Col-0 wild type leads to a similar cell wall composition as periodic starvation in rosettes of starch-deficient *pgm* mutants. Galactose content is significantly but reversibly reduced upon XN treatment, indicating dynamic remodeling of cell wall composition.

## Starvation-induced beta-galactosidases release galactose from the cell wall

In the light of the known alterations of RG-I side chains in *pgm* (Engelsdorf *et al*., 2017), the starvation-dependent reduction in cell wall galactose suggested that β-1, 4-galactan might be remobilized upon carbohydrate shortage. Transcriptomics data by Usadel *et al*. (2008) indicated that the 𝛽-galactosidases *BGAL1, 2, 4, 6* and *10* are transcriptionally induced in *pgm* during the dark phase (*BGAL1, 2, 4*) and/or after XN (*BGAL1, 2, 4, 6, 10*) (Table S1). Using qRT-PCR, we investigated the induction of these five *BGAL* genes after 24 hours of XN, when galactose release was around 50% (cf. Fig. 1a). The results confirmed a significant upregulation of *BGAL1, 2, 4* and *10* after XN, while *BGAL6* transcript was not significantly different between XN and control conditions (Fig. 2a). *BGAL1, BGAL4* and *BGAL10* exhibited a more than ten-fold induction after XN compared to EN (Fig. 2b). To elucidate the contributions of these three *BGAL* genes to the XN-induced reduction in cell wall galactose, we examined two independent T-DNA insertion lines per gene. We identified *bgal1-1, bgal1-2, bgal4-1* and *bgal4-2* as loss-of-function mutants (Fig. S3), and obtained *bgal10-1* and *bgal10-2*, which have been previously described as xyloglucan 𝛽-galactosidase mutants (Sampedro et al., 2012). An analysis of the galactose content in the rosette cell walls of these mutants after 72 hours XN indicated that the decrease in galactose was less pronounced in *bgal1-1* (16,1%)*, bgal1-2* (16,9%)*, bgal4-1* (14,5%), and *bgal4-2* (13,2%) compared to Col-0 (20,3%), while in *bgal10-1* (22,9%) and *bgal10-2* (20,2%) XN-induced galactose release from the cell wall was not impaired (Fig. 2c).

**Figure 2.**
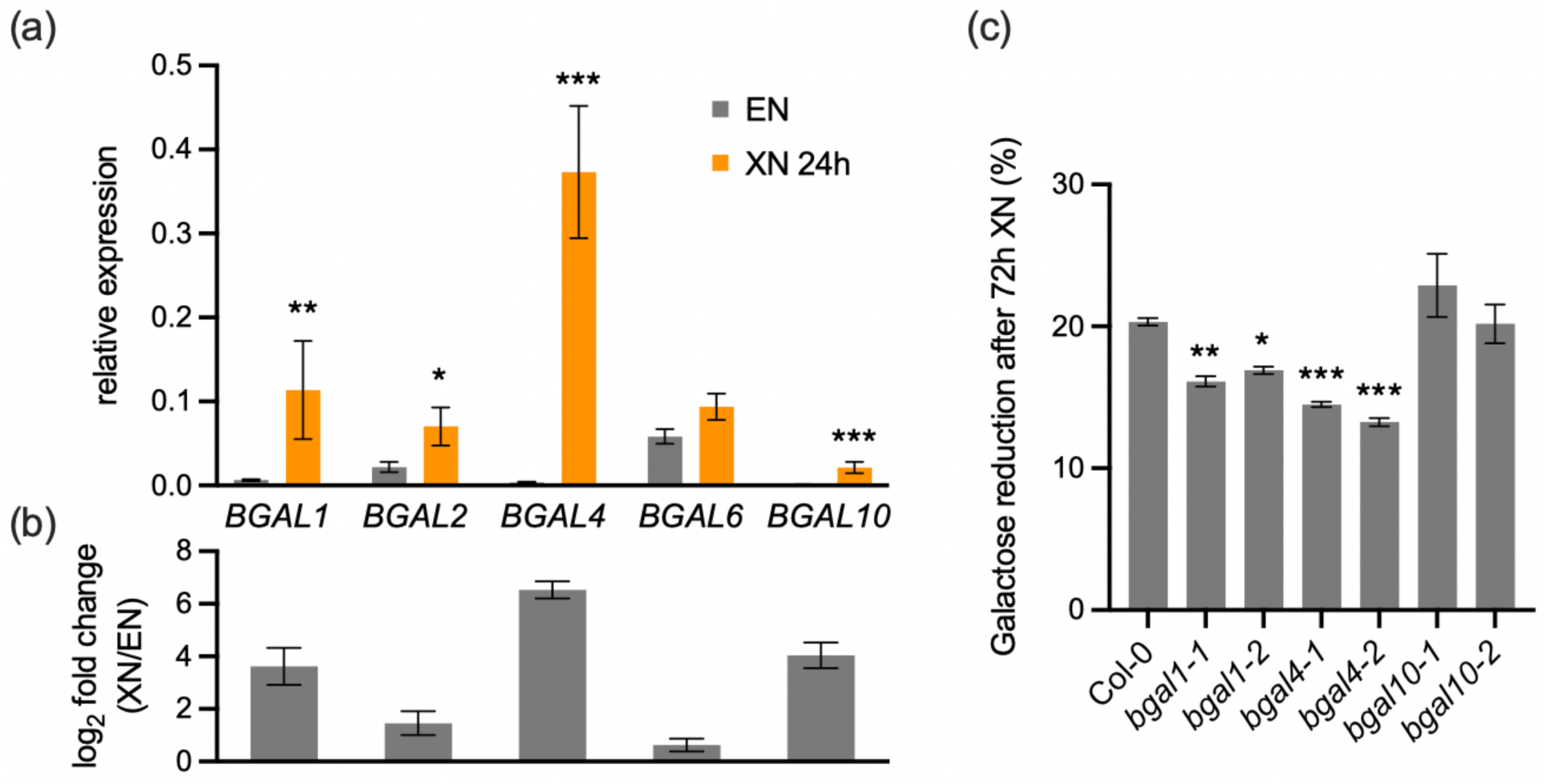
: *BGAL1* and *BGAL4* contribute to starvation-induced release of cell wall galactose. (a) The relative expression of *BGAL1*, *BGAL2*, *BGAL4*, *BGAL6* and *BGAL10* was determined by qRT-PCR in Col-0 rosettes at the end of the night (EN) and after 24h of extended night (XN). Values are means (n=4) and error bars represent the SEM. Asterisks indicate statistically significant diXerences between XN 24h and EN according to a Student’s t-Test (*p<0.05, **p<0.01, ***p<0.001). (b) Bars represent the log2 fold change of values depicted in (a) +/- SEM. (c) The reduction of galactose in cell walls of Col-0, *bgal1-1*, *bgal1-2*, *bgal4- 1*, *bgal4-2*, *bgal10-1* and *bgal10-2* rosettes was quantified after 72h XN relative to EN. Values are means (n=4-5) and error bars represent the SEM. Asterisks indicate statistically significant diXerences to Col-0 according to one-way ANOVA and Dunnett’s multiple comparisons test (*p<0.05, **p<0.01, ***p<0.001).

These results indicate that BGAL1 and BGAL4 contribute to starvation-dependent galactan degradation, while BGAL10 is not required for the release of galactose from leaf cell walls during XN.

### Galactose salvage is induced upon starvation and cannot be prevented in *uge1 uge3* mutants

Published transcriptomics data indicated an induction of the galactan salvage pathway and a simultaneous attenuation of galactan synthesis after XN in the wild type and in *pgm* at EN (Usadel *et al*., 2008, Engelsdorf *et al*., 2017). We hypothesized that galactose salvage might be important to provide sugars to rosette leaves under starvation, while at the same time a reduced biosynthesis of galactan might impair the cell wall structure and penetration resistance. To further characterize the transcriptional activation of the salvage pathway, we investigated the expression of *UGE1* and *UGE3*, which have been proposed to act in carbohydrate catabolism (Barber *et al*., 2006), via qRT-PCR. In both Col-0 and *pgm*, XN lead to a significant increase in *UGE1* expression (Fig. 3a). *UGE3* transcripts showed a similar pattern but had a lower expression level than *UGE1* (Fig. 3b). Since galactose released from the cell wall via BGAL activity has to pass through the salvage pathway for re-utilization by the plant, this data suggested that the metabolic flux through the salvage pathway might be increased compared to carbon replete wild type plants under control conditions. To investigate if loss of UGE1 and UGE3 causes a backlog of UDP-galactose upon starvation, we crossed *pgm* with *uge1 uge3* double mutants and investigated nucleotide sugar levels at EN and after XN. Surprisingly, UDP-galactose amount was neither increased in *uge1 uge3* rosettes compared to the wild type upon starvation, nor in *pgm uge1 uge3* rosettes compared to *pgm*, indicating that UGE1 and UGE3 are not the only UGE isoforms responsible for galactose salvage (Fig. 3c). Instead, the amounts of UDP-galactose and UDP-glucose were strongly reduced after XN and in *pgm*, probably reflecting reduced overall carbohydrate availability (Fig. 3c,d). The cell wall composition of *uge1 uge 3* after 72 hours of XN was not altered compared to Col-0, which excludes the possibility that recycled UDP-galactose is immediately reused for cell wall synthesis (Fig. S4). To investigate if impaired *UGE1* and *UGE3* expression influences susceptibility of *pgm* to *C. higginsianum*, we infected Col-0, *uge1 uge3*, *pgm*, and *pgm uge1 uge3* rosettes with the fungus and assessed fungal genomic DNA as a proxy for fungal biomass at 4 days post infection (dpi) via qPCR. Susceptibility to *C. higginsianum* was not affected by the loss of *UGE1* and *UGE3* (Fig. 3e).

**Figure 3.**
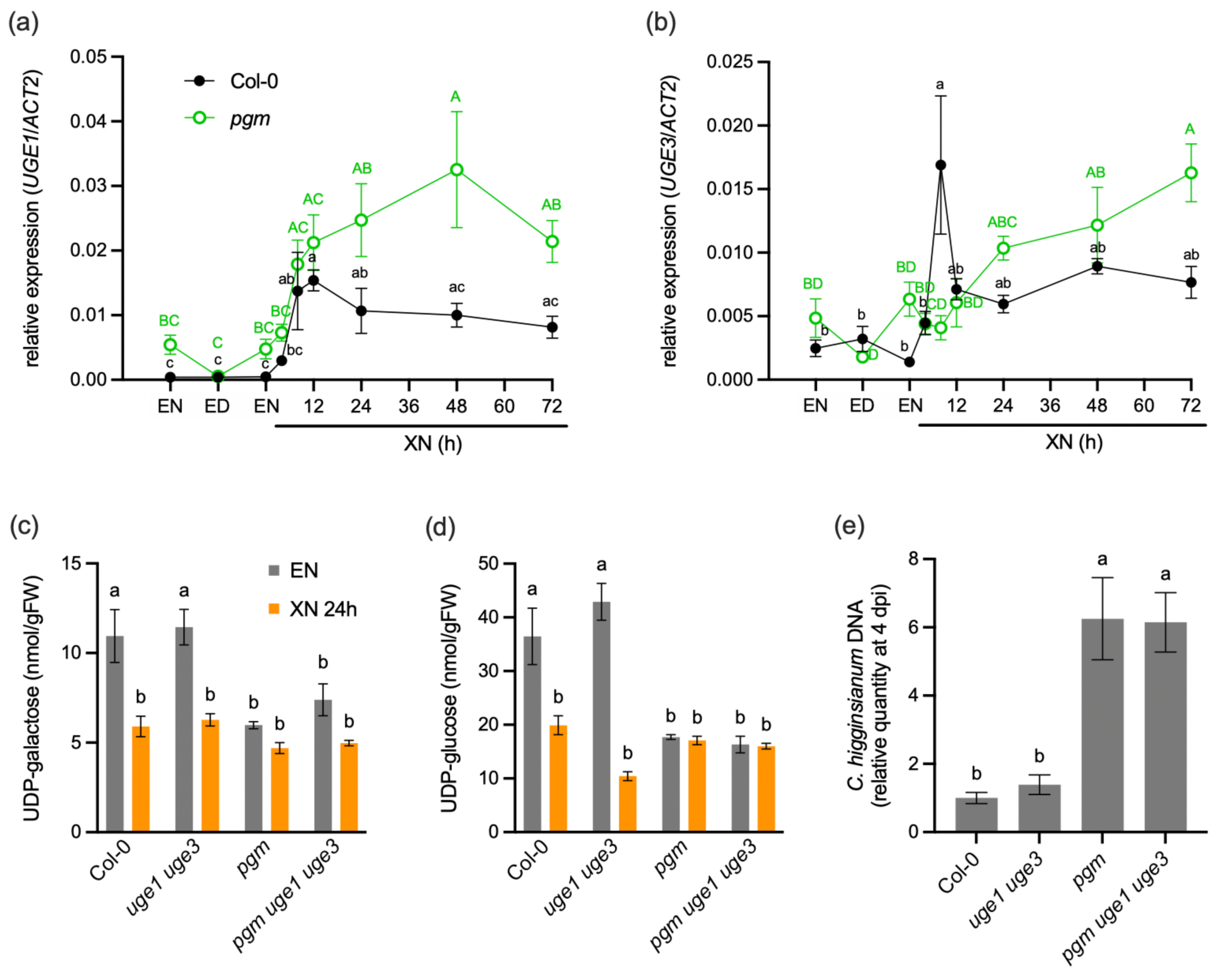
: Starvation-induced *UGE1* and *UGE3* expression is dispensable for galactose salvage. The relative expression of (a) *UGE1* and (b) *UGE3* was determined in rosette leaves of Col-0 (black) and *pgm* (green) via qRT-PCR analysis at the end of the night (EN) and the end of the day (ED) in a 12h light / 12h dark cycle, as well as after an extended night (XN) of 4, 8, 12, 24, 48 and 72h. Values are means (n=3-4) and error bars represent the SEM. DiXerent letters indicate statistically significant diXerences according to one-way ANOVA and Tukey’s multiple comparisons test for Col-0 (black) and *pgm* (green, capital letters) samples, respectively (α = 0.05). (c) UDP-galactose and (d) UDP-glucose was quantified in Col-0, *uge1 uge3*, *pgm* and *pgm uge1 uge3* rosettes at EN and 24h XN. Values are means (n=5) and error bars represent the SEM. DiXerent letters indicate statistically significant diXerences according to two-way ANOVA and Tukey’s multiple comparisons test (α = 0.05). (e) Rosettes of the indicated genotypes were spray-infected with *Colletotrichum higginsianum* and the fungal proliferation assessed via quantification of fungal DNA via qPCR at 4 days post infection (dpi). Values are means (n=6) and error bars represent the SEM. DiXerent letters indicate statistically significant diXerences according to one-way ANOVA and Tukey’s multiple comparisons test (α = 0.05).

To summarize, transcriptional activation of *UGE1* and *UGE3* indicated increased galactose salvage upon XN and in *pgm*. However, loss of *UGE1* and *UGE3* is neither sufficient to cause UDP-galactose backlog nor to alter starvation-dependent pathogen susceptibility, indicating that they do not represent the main catabolic UGE isoforms.

## Increased cell wall galactan limits starvation-dependent pathogen entry

The reduction in cell wall galactose within hours after onset of starvation, together with attenuated *GALS1* expression, indicated that galactan synthesis might be reduced in addition to the mobilization of galactose from existing galactan chains (Usadel *et al*., 2008, Engelsdorf *et al*., 2017). A qRT-PCR time course analysis of *GALS1* confirmed increased expression at the end of the day (ED) and strongly reduced *GALS1* transcript amounts at EN and after XN (Fig. 4a). *GALS1* transcripts were depleted in *pgm* at EN and in Col-0 after 72 hours XN (Fig. 4a).

**Figure 4.**
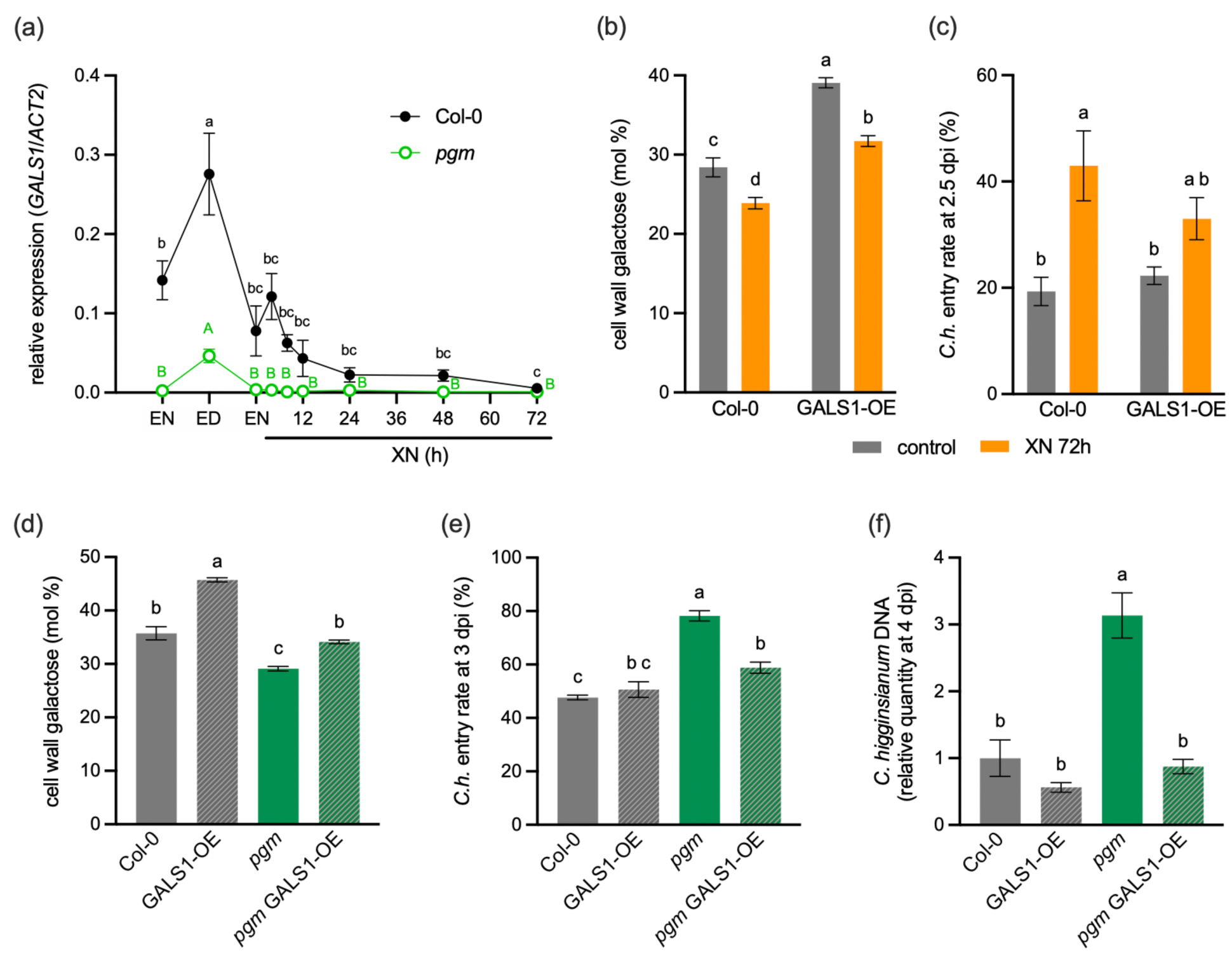
: Galactan content in leaf cell walls contributes to resistance to *Colletotrichum higginsianum infection.* (a) The relative expression of *GALS1* was determined in rosette leaves of Col-0 (black) and *pgm* (green) via qRT-PCR analysis at the end of the night (EN) and the end of the day (ED) in a 12h light / 12h dark cycle, as well as after 4, 8, 12, 24, 48 and 72h extended night (XN). Values are means (n=3-4) and error bars represent the SEM. DiXerent letters indicate statistically significant diXerences according to one-way ANOVA and Tukey’s multiple comparisons test for Col-0 (black) and *pgm* (green, capital letters) samples, respectively (α = 0.05). (b) Molar percentage of galactose in cell wall neutral monosaccharides and (c) entry rate of *Colletotrichum higginsianum* at 2.5 days post infection (dpi) in Col-0 and GALS1-OE under control conditions (EN) and after 72h XN. Values are means (b, n=4; c, n=5-7) and error bars represent the SEM. DiXerent letters indicate statistically significant diXerences according to two-way ANOVA and Tukey’s multiple comparisons test (α = 0.05). (d) Molar percentage of galactose in cell wall neutral monosaccharides, (e) *C. higginsianum* entry rate at 3 dpi and (f) *C. higginsianum* DNA at 4 dpi were quantified in Col-0, GALS1-OE, *pgm*, and *pgm* GALS1-OE. Values are means (d, n=4; e and f, n=5) and error bars represent the SEM. DiXerent letters indicate statistically significant diXerences according to one-way ANOVA and Tukey’s multiple comparisons test (α = 0.05).

*GALS1* overexpressing lines (GALS1-OE) exhibit about 200-fold increased *GALS1* expression and high β-1, 4-galactan levels (Fig. S5; Liwanag *et al*., 2012). We employed these GALS1-OE lines to test the hypothesis that an increased galactan content in rosette cell walls might protect against starvation-dependent cell wall defects and associated susceptibility towards *C. higginsianum*. We confirmed that GALS1 overexpression was not affected by 72 hours of XN (Fig. S5a). GALS1-OE cell walls showed intact galactose release after XN, but galactose amounts remained significantly higher than in non-starved Col-0 rosettes (Fig. 4b). To test if the higher galactose content influences penetration resistance to *C. higginsianum*, we determined the fungal entry rate. After 72 hours XN compared to control conditions, fungal entry was significantly increased in Col-0, but not in GALS1-OE, suggesting that an increased cell wall galactan content mitigates the penetration success of the fungus (Fig. 4c). As the cell wall composition of *pgm* rosettes resembles that of Col-0 after XN, we hypothesized that *GALS1* overexpression would also reduce penetration success of *C. higginsianum* in the *pgm* background. We crossed *pgm* with GALS1-OE plants and confirmed that *GALS1* expression and galactose content were increased in the resulting line (Fig. 4d, Fig. S5b). In both Col-0 and *pgm* rosettes, *GALS1* overexpression led to an altered composition of neutral cell wall monosaccharides, with an increased molar ratio of galactose, whereas the major pectic sugar galacturonic acid was not affected (Fig. S6). The fungal entry rate in *pgm* GALS1-OE was significantly reduced compared to *pgm*, indicating increased penetration resistance (Fig. 4e). Galactan-dependent resistance of *pgm* to *C. higginsianum* was also detected via qPCR of fungal DNA at 4 dpi, demonstrating that the reduced entry rate leads to reduced fungal proliferation and a suppression of *pgm* hypersusceptibility (Fig. 4f).

Collectively, our data show that carbohydrate starvation causes the release of galactose from pectic β- 1, 4-galactan via BGAL1 and BGAL4 activity, while reduced *GALS1* expression prevents the flux of carbon into galactan biosynthesis (Fig. 5). The resulting alteration in cell wall composition critically affects penetration resistance against *C. higginsianum*, which can be rescued by increased galactan synthesis.

**Figure 5.**
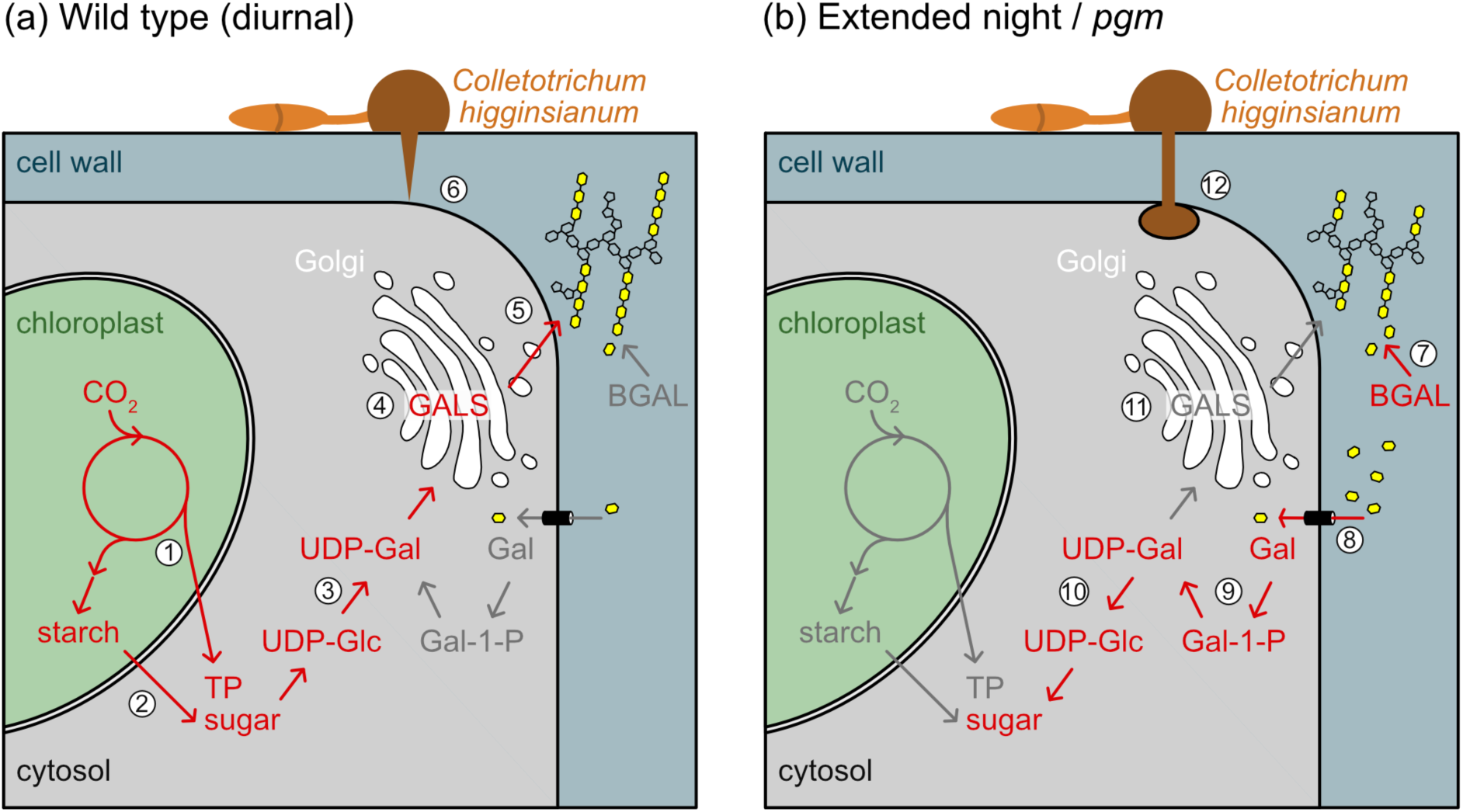
: Model illustrating cell wall galactan remodeling upon starvation and its impact on fungal penetration resistance. (a) Carbon usage for galactan synthesis and penetration resistance in Arabidopsis wild type rosettes during a regular diurnal cycle: (1) Carbon fixed in the Calvin cycle during the day is exported to the cytosol as triose phosphate (TP) and stored in the form of starch as a reservoir for the night. (2) Starch is degraded to maltose and glucose during the night and utilized for UDP-glucose (UDP-Glc) synthesis. (3) UDP-galactose (UDP-Gal) is formed from UDP-Glc via the action of UDP-Glc 4-epimerases (UGEs). (4) After import into the Golgi, UDP-Gal serves as substrate for β-1,4-galactan synthesis by GALACTAN SYNTHASE (GALS). (5) Galactan is secreted into the apoplast and linked to pectic rhamnogalacturonan-I. (6) An intact cell wall limits penetration by the hemibiotrophic fungus *Colletotrichum higginsianum*. (b) Cell wall galactan recycling during an extended night or at the end of the night in starch-deficient *phosphoglucomutase* (*pgm*) mutants: (7) BETA-GALACTOSIDASE (BGAL) expression is induced upon starvation and leads to the release of Gal from galactan chains. (8) Gal is imported to the cytosol via sugar transporters. (9) The Gal salvage pathway converts Gal to Gal-1-P and UDP-Gal. (10) UGEs epimerize UDP-Gal to UDP-Glc, which can be utilized for cellular metabolism. (11) GALS1 expression is strongly downregulated, contributing to a decreasing galactan amount in the cell wall and (12) impaired penetration resistance against *C. higginsianum*.

## Discussion

Structural carbohydrates of the cell wall represent the largest carbon sink in plants (Barnes and Anderson, 2018). To date it is not clear to what degree carbon can be remobilized from cell walls, and which cell wall polymers would be prioritized for recycling. Previous reports indicated that dark-induced starvation leads to an induction of glycoside hydrolases and an altered cell wall composition (Lee *et al*., 2007, Poschet *et al*., 2010). In Arabidopsis mutants with impaired starch synthesis or mobilization, cell wall arabinose and galactose levels were reduced compared to the wild type, indicating that diurnal carbohydrate shortage in these lines influences polysaccharide biosynthesis and/or degradation (Engelsdorf *et al*., 2017).

Here, we undertook a detailed comparison of the cell wall monosaccharide composition in rosettes of wild type and the starch-deficient *pgm* mutant upon dark-induced starvation. Our data reveal that galactose is the major sugar released from cell walls under these conditions. Of note, the galactose levels in *pgm* cell walls under control conditions resembled that of the wild type after two days of starvation. This is in line with previously published transcript and metabolite datasets showing similarities between carbon-limited *pgm* and XN starved wild type leaves (Gibon *et al*., 2006, Usadel *et al*., 2008).

In *pgm,* galactose content was not further reduced after 2 days of XN, indicating that it cannot be reduced below ∼25 % of the total neutral cell wall monosaccharides. Indeed, the *gals1/2/3* triple mutant lacking all three GALS isoforms has a similar galactose content as the starved *pgm* mutant. As no additional galactose is released from the cell walls of *gals1/2/3* mutants upon XN, we speculate that carbon depletion induces degradation of galactan side chains in RG-I. Arabinan side chains in RG-I seem to be less affected by dark-induced starvation. While arabinose is also reduced in cell walls of starch-deficient mutants (Engelsdorf *et al*., 2017), it was only mildly altered up to 48 hours XN in wild type rosettes. This data indicates that the low arabinose content in the *pgm* mutant is less connected to starvation. Altered galactan content in *gals1/2/3* and GALS1-OE lines does not cause obvious visible phenotypes (Ebert *et al*., 2018), suggesting that it may represent a suitable and flexible source of carbon to be recycled from the cell wall pool. Consistent with this model, we found that the galactose levels in wild type cell walls can be almost fully recovered after three days of returning starved plants into a regular light/dark cycle.

We identified four beta-galactosidase genes that were significantly upregulated after 24 hours of XN. Mutant analysis demonstrated that at least two of them, *BGAL1* and *BGAL4*, contribute to the decline of galactose in leaf cell walls during XN. This finding corroborates a previous report that BGAL4 is secreted upon carbohydrate starvation in detached leaves (Lee *et al*., 2007). Seven additional BGAL isoforms have been reported to be expressed in leaves (Ahn *et al*., 2007), of which *BGAL2*, *BGAL6* and *BGAL10* may be upregulated after starvation (Usadel *et al*., 2008, Poschet *et al*., 2010). BGAL10 has been reported to represent the main BGAL isoform acting on xyloglucan (Sampedro *et al*., 2012). As we did not observe a reduced release of galactose in *bgal10* mutants after XN, our data indicate that galactose released upon starvation does not originate from xyloglucan. This is in line with the GALS- dependency of the released galactose pool and suggests that BGAL1 and BGAL4 degrade RG-I galactan side chains upon XN treatment.

Released galactose is likely taken up from the cell wall by monosaccharide transporters of the STP family (Poschet *et al*., 2010, Barnes and Anderson, 2018). Galactose can be recycled for cellular metabolism via a salvage pathway, which involves phosphorylation via GALK and conversion into UDP-galactose by USP (Reiter, 2008, Egert *et al*., 2012, Geserick and Tenhaken, 2013). Arabidopsis encodes five UGE isoforms that are able to interconvert UDP-galactose and UDP-glucose. In line with the previously made suggestion that the UGE1 and UGE3 isoforms are mainly acting in carbohydrate catabolism (Barber *et al*., 2006), we detected a strong upregulation of *UGE1* and *UGE3* upon nocturnal starvation in *pgm* and upon XN, while none of the other *UGE* isoforms was differentially expressed under both conditions (Usadel *et al*., 2008, Engelsdorf *et al*., 2017). However, loss of *UGE1* and *UGE3* did not cause differences in UDP-galactose or UDP-glucose levels in wild type or *pgm* backgrounds at EN or after XN, which suggests that they can be functionally replaced by the remaining three UGE isoforms. We refrained from further analysis of higher order UGE mutants, as those have been reported to have extreme dwarf phenotypes (Rösti *et al*., 2007). Nevertheless, it will be exciting to further explore the specific functions of the individual UGE isoforms in the future.

In addition to the induction of BGAL expression and of the galactose salvage pathway, XN treatment led to the downregulation of the galactan synthase genes *GALS1*, *GALS2* and *GALS3* (Usadel *et al*., 2008, Engelsdorf *et al*., 2017). All three GALS isoforms contribute to galactan synthesis *in vivo*, with GALS1 and 3 being the main isoforms expressed in leaves and loss of GALS1 having the strongest effect on galactose content in leaf cell walls (Liwanag *et al*., 2012, Ebert *et al*., 2018). Of note, *GALS1* expression is close to the detection limit in *pgm* leaves already at EN, indicating that galactan synthesis is halted in *pgm* every night when soluble sugar reserves are depleted. The gene expression patterns of *GALS1/2/3*, *UGE1/3* and *BGAL1/4* together indicate that galactan synthesis and degradation are inversely regulated and balanced depending on the available carbohydrate budget. Our investigation of GALS1-OE lines showed that the relative amount of galactose present in cell walls of rosette leaves can be increased by more than 30% and remains higher in GALS1-OE than in Col-0 (at EN) even after 72 hours XN. Nevertheless, the additional galactose in GALS1-OE cell walls is amenable to starvation-induced release. By contrast, reduced galactose levels in *gals1/2/3* triple mutants were not further reduced upon XN, indicating that only RG-I galactan side chains are recycled upon starvation. Based on our findings, we propose that the galactan side chains in RG-I represent a dynamic carbohydrate reserve in leaf cell walls that can be utilized upon starvation.

The cell wall represents a major penetration barrier to fungal infection and infection success often depends on modifications caused by both, host plant and pathogen (Munzert and Engelsdorf, 2025). We have previously reported that periodic nocturnal starvation in starch-free mutants leads to reduced arabinose and galactose content and reduced detection of RG-I/arabinogalactan epitopes in rosette leaf cell walls (Engelsdorf *et al*., 2017). Comparative infection experiments with other mutants defective in cell wall composition indicated that RG-I composition (in *mur8*; Mertz *et al*., 2012) and total pectin abundance (in *pmr5 pmr6*; Vogel *et al*., 2004) are critical for penetration resistance against *C. higginsianum* (Engelsdorf *et al*., 2017). In contrast, specific impairment of arabinose (*arad1, mur4*) and galactose (*gals1*) content in cell walls did not influence pathogen susceptibility during a regular diurnal cycle (Engelsdorf *et al*., 2017). Here, we show that galactose release from rosette cell walls critically affects penetration resistance in plants that suffer from periodic (*pgm*) or dark-induced (XN) starvation. While sugars derived from photosynthetic carbon assimilation can be utilized for galactan synthesis throughout a regular diurnal cycle (Fig. 5a), starvation causes a pause of sugar supply from the chloroplast. Under these conditions, BGALs, STPs, UGEs are induced to initiate recycling of cell wall galactose via the galactose salvage pathway (Fig. 5b, Usadel *et al*., 2008). As a consequence of starvation-dependent alterations in cell wall composition, penetration resistance against *C. higginsianum* is impaired, leading to an increased fungal entry rate. Our data reveal that overexpression of GALS1, counteracting the starvation-induced reduction of cell wall galactose, is sufficient to suppress the increased fungal entry rates upon starvation and the hypersusceptibility of *pgm*.

In summary, we present evidence that galactose is recycled from RG-I side chains in Arabidopsis leaf cell walls during carbohydrate shortage enabling the plant to tap into cell wall sugar reserves. Due to the highly dynamic organization of pectic polymers (Anderson and Pelloux, 2025), they might be a preferred substrate for sugar recycling. However, the composition of pectin is also of crucial importance for fungal penetration resistance, which is consequently compromised under starvation conditions. This tradeoff between sugar supply for plant metabolism and preformed pathogen defense highlights the importance of cell wall surveillance to sense and maintain functional integrity under stress conditions.

## Materials and Methods

### Plant material and growth conditions

*Arabidopsis thaliana* plants were grown in a 12 h light (22°C) / 12 h dark (20°C) cycle at a photon flux density of 110 µmol m^-2^ s^-1^. Genotypes used in this study are listed in Supplementary Table S2. Five days prior to an infection with *Colletotrichum higginsianum* or extended night, plants were fertilized with 40 ml 0.1% Wuxal Super (Aglukon) fertilizer per pot. 5 week-old plants were used for starvation and infection experiments.

### *Colletotrichum higginsianum* infection assays

*Colletotrichum higginsianum* isolate MAFF 305635 (Ministry of Forestry and Fisheries, Japan) was grown on oatmeal agar plates (5% [w/v] shredded oatmeal, 1.2% [w/v] agar) for 7 days in a 16 h light (22°C) / 8 h dark (20°C) cycle at a photon flux density of 110 µmol m^-2^ s^-1^. Conidia were harvested by rinsing plates with deionized water, the conidia titer was adjusted to 2 · 10^6^ conidia ml^-1^ and the suspension was immediately used for infection experiments. Spray infection was performed as described by Voll *et al*. (2012). Briefly, inoculation was performed at the end of the light phase and high humidity maintained until 2.5 days post infection to provide consistent conditions for humidity-dependent fungal penetration. Fungal entry rates were determined under the microscope after staining fungal structures with trypan blue as described (Engelsdorf *et al*., 2017). Quantification of the relative fungal DNA content in infected leaves was performed by qPCR analysis of *Ch*TrpC as previously described (Engelsdorf *et al*., 2013).

### Gene expression analysis

Total RNA was isolated using a NucleoSpin Plant and Fungi RNA Mini Kit (Macherey-Nagel). One microgram of total RNA was treated with RNase-Free DNase (Thermo Scientific) and complementary DNA (cDNA) synthesized with a RevertUP II Reverse Transcriptase Kit (Biotechrabbit). Quantitative reverse transcription polymerase chain reaction was performed using Blue S’Green (Biozym) or PowerUp (Applied Biosystems) SYBR green mix and a Bio-Rad CFX Connect RT-PCR system. Primers are listed in Supplementary Table S3 and were diluted according to the manufacturer’s specifications. *ACT2* was used a reference in all experiments.

### Quantification of soluble sugars and starch

Soluble sugars and starch were analyzed as described by Voll *et al*. (2003). Briefly, snap-frozen leaves were extracted twice with 80% ethanol at 80°C. After evaporation of ethanol and resuspending in deionized water, soluble glucose, fructose and sucrose were quantified in a coupled enzymatic assay using a Tecan infinite 200Pro microtiter plate reader. Extracted leaves were ground in 0.2 M KOH and starch was solubilized by heating for 45 min @ 95°C. After degradation by a mixture of α-amylase and amyloglucosidase, resulting glucose was quantified as described above.

### UDP-galactose and UDP-glucose analysis

The extraction of UDP-galactose and UDP-glucose was done as described previously (Rautengarten *et al*., 2019). Briefly, whole Arabidopsis rosettes were ground in liquid nitrogen and extracted in ice-cold chloroform / methanol (3:7). After incubation for 2 h at -20°C, ice-cold water was added, and the upper phase was collected after 5 min centrifugation at 4°C and 30,000 x g. Extraction was repeated two times and combined extracts were lyophilized. 10 mM ammonium bicarbonate was added to the dried sample, mixed, and briefly spun down. Equilibration of an EnviCarb SPE (Sigma-Aldrich) was performed using 80% acetonitrile with 0.1% trifluoroacetic acid (TFA) followed by 2 mL of water. The sample in 10 mM ammonium bicarbonate was added to the SPE column, and the column washed with water, 25% acetonitrile and finally 50 mM triethylamine/acetic acid (TEAA) pH 7.0. Elution was done with 25% acetonitrile with 50 mM TEAA pH 7.0. Following an overnight lyophilization step, the UDP-sugars were resuspended in ice-cold water and analysed by LC-MS/MS. LC-MS/MS was performed using porous graphitic carbon as the stationary phase on an 1100 series HPLC system (Agilent Technologies) and a 4000 QTRAP LC/MS/MS system (SCIEX) equipped with a TurboIonSpray ion source using methods previously described (Rautengarten *et al*., 2014).

### Cell wall analysis

Whole Arabidopsis rosettes were ground in liquid nitrogen and extracted three times with 80% ethanol at 80°C and washed with deionized water. Alcohol-insoluble residue (AIR) was dried by lyophilization, weighed in 2-mL tubes with screw caps and used for cell wall hydrolysis as described by Yeats *et al*. (2016). Briefly, neutral cell wall sugars and uronic acids from non-crystalline cell wall matrix polymers were hydrolyzed in 4% (w/v) sulfuric acid by autoclaving at 121°C for 60 min. Monosaccharides were diluted with ultrapure water and ribose was added as an internal standard. Analysis was performed via high-performance anion-exchange chromatography with pulsed amperometric detection (HPAEC-PAD) using a biocompatible Knauer Azura HPLC system and an Antec Decade Elite SenCell detector heated to 40°C. Monosaccharides were separated on a Thermo Fisher Dionex CarboPac PA20 BioLC analytical column (3 x 150 mm) equipped with a CarboPac PA20 BioLC guard column (3 x 30 mm) using the following solvent gradient of (B) 10 mM NaOH and (C) 700 mM NaOH in (A) water, at 0.4 mL /min flow rate: 0 to 25 min: 20% B; 25 to 28 min: 20% to 0% B, 0% to 70% C; 28 to 33 min: 70% C; 33 to 35 min: 70% to 100% C; 35 to 38 min: 100% C; 38 to 42 min: 0% to 20% B, 100% to 0% C; 42 to 60 min: 20% B.

## Statistical analysis

Statistical analysis was performed with GraphPad Prism (Version 10.4.0). Specific statistical methods, sample size, significance level and p-values are indicated in the respective figure legends.

## Acknowledgements

This work was funded by the Deutsche Forschungsgemeinschaft (DFG, German Research Foundation) grant EN 1071/3-1 to T.E. and B.E. would like to acknowledge funding from the RUB Research School, the University of Melbourne Botany Foundation, and Australian Research Council grant DP220101544. We would like to thank Georg Seifert (BOKU Vienna) for providing *uge1-1 uge3-2* seeds, Henrik Vibe Scheller (UC Berkeley) for GALS1-OE seeds and Lars Voll (Philipps-Universität Marburg) for critical reading of the manuscript.

## Author contributions

Conceptualization: TE; Investigation: LR, RNM, KSM-E, ST, CR, VT, KLH, LZ, TE; Funding acquisition: BE, TE; Supervision: RNM, KSM-E, BE, TE; Writing – original draft: TE; Writing – review & editing: LR, RNM, KSM-E, ST, CR, VT, KLH, LZ, BE, TE.

---

**Figure S1.**
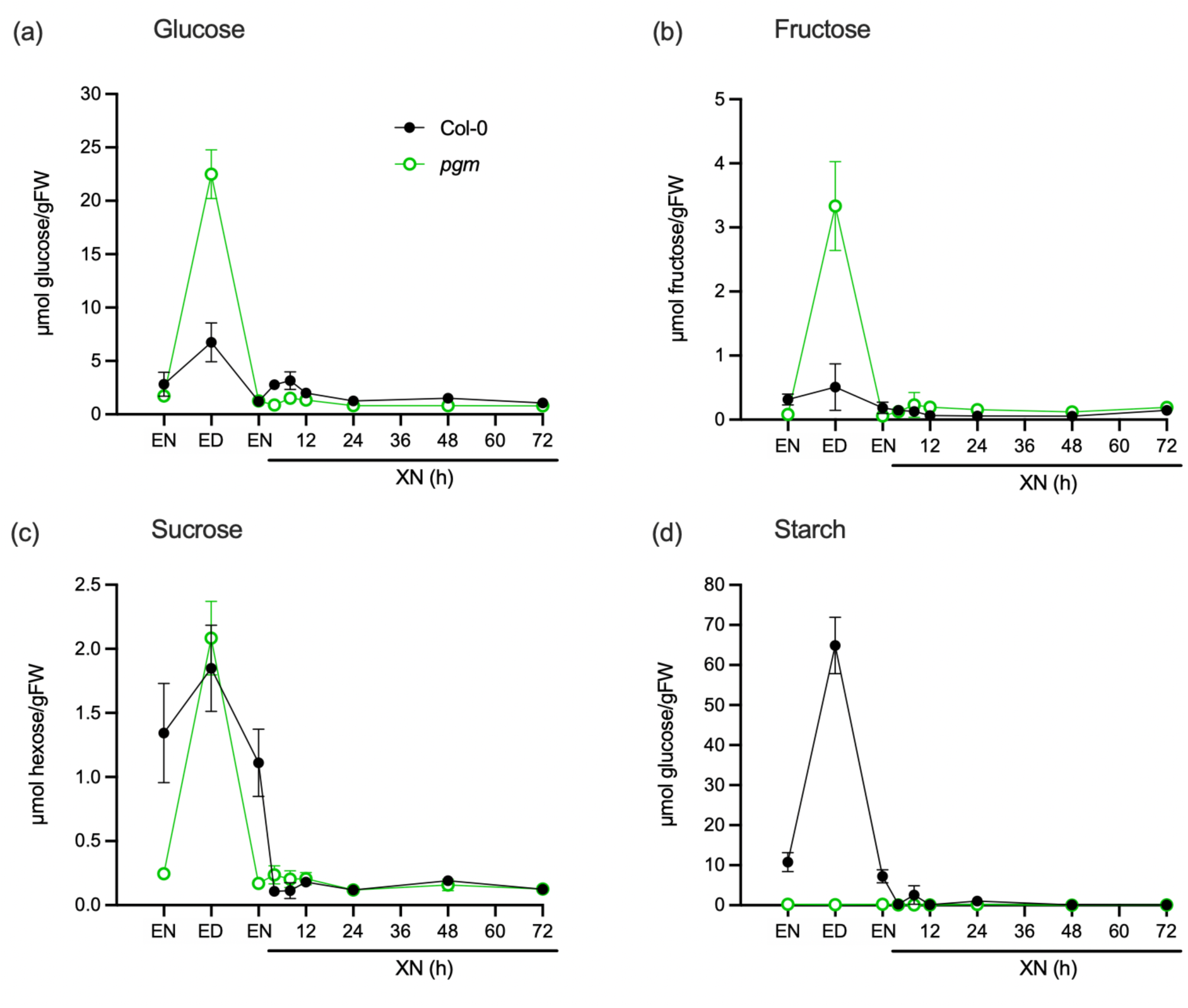
: **Carbohydrate depletion during dark-induced starvation in leaves.** (a) Glucose, (b) fructose, (c) sucrose and (d) starch were quantified in rosette leaves of Col-0 (black) and *pgm* (green) at the end of the night (EN) and the end of the day (ED) in a 12h light / 12h dark cycle, as well as after 4, 8, 12, 24, 48 and 72h extended night (XN). Values are means (n=4) and error bars represent the SEM.

**Figure S2.**
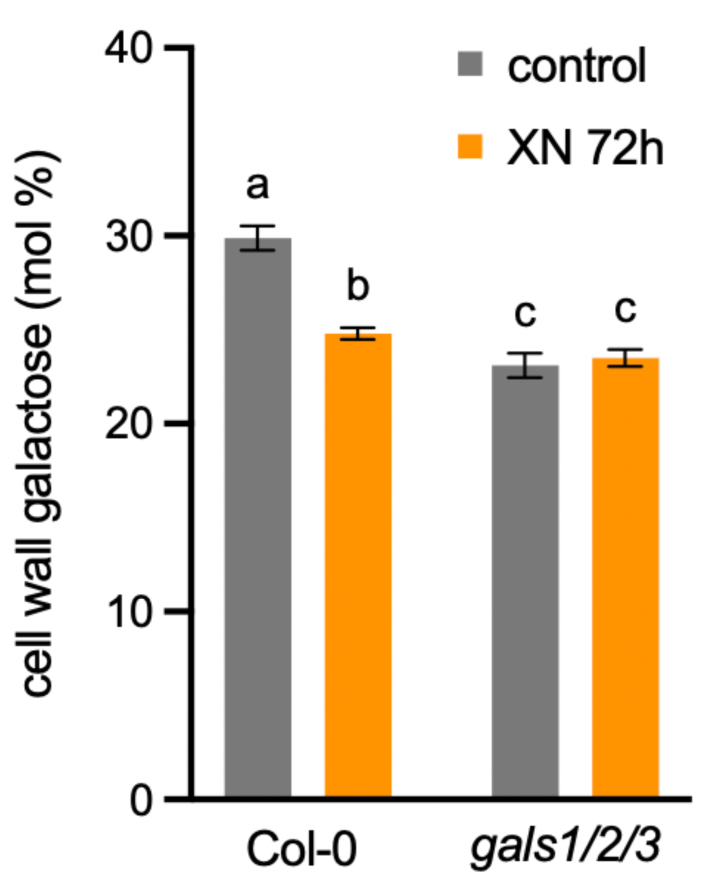
: Galactose content in *gals1/2/3* mutants. Molar percentage of galactose in cell wall neutral monosaccharides in Col-0 and *gals1/2/3* under control conditions (EN) and after 72h XN. Values are means (n=3-4) and error bars represent the SEM. DiXerent letters indicate statistically significant diXerences according to two-way ANOVA and Tukey’s multiple comparisons test (α = 0.05).

**Figure S3:**
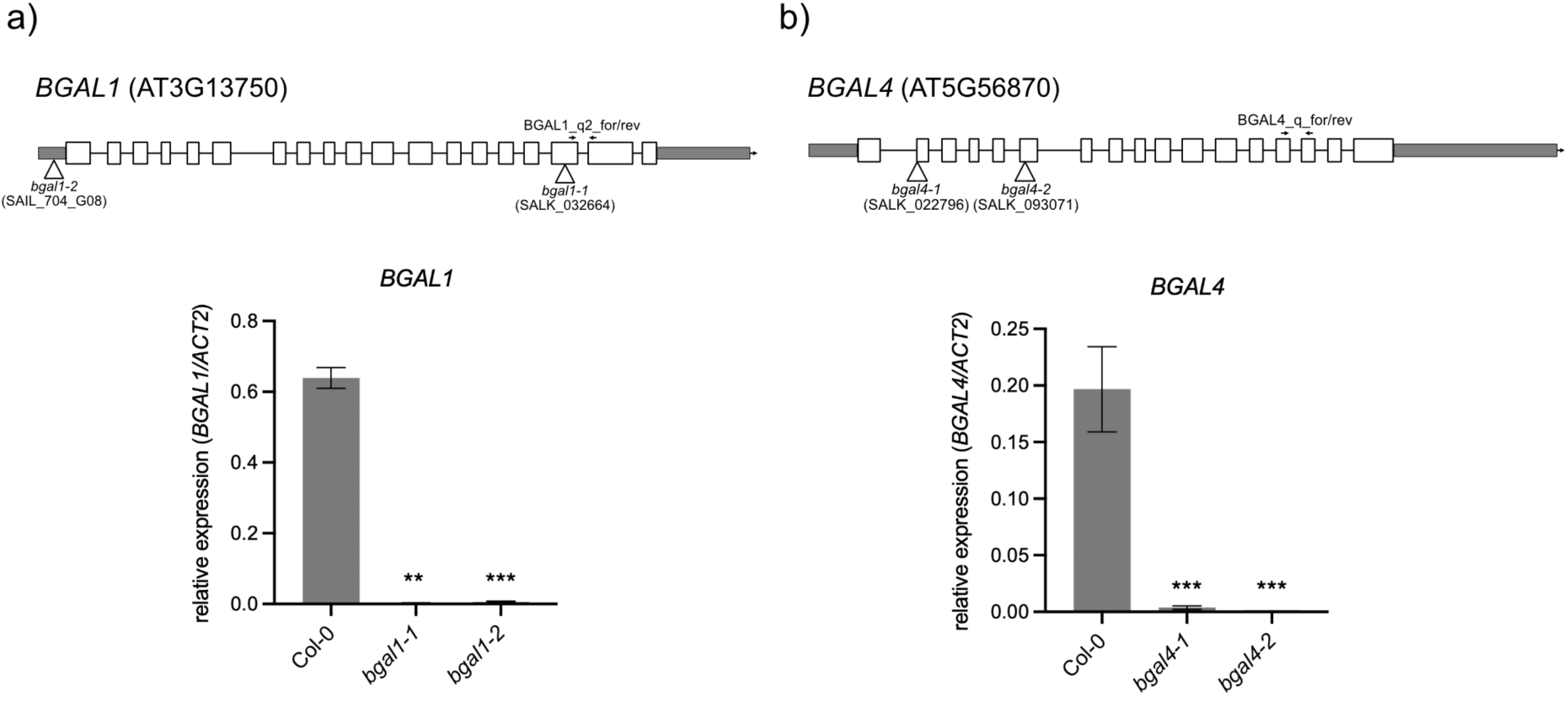
Characterization of *bgal1* and *bgal4* mutants. The relative expression of (a) *BGAL1* in Col-0, *bgal1-1* and *bgal1-2* rosettes and (b) *BGAL4* in Col-0, *bgal4-1* and *bgal4-2* rosettes were determined by qRT-PCR after 24h of extended night. Sketches of the genes indicate exons (white boxes), introns (lines), UTR regions (grey boxes), T-DNA insertion sites (triangles), and primer binding sites (arrows). Values are means (n=4) and error bars represent the SEM. Asterisks indicate statistically significant diXerences to Col-0 according to a Student’s t-Test (**p<0.01, ***p<0.001).

**Figure S4:**
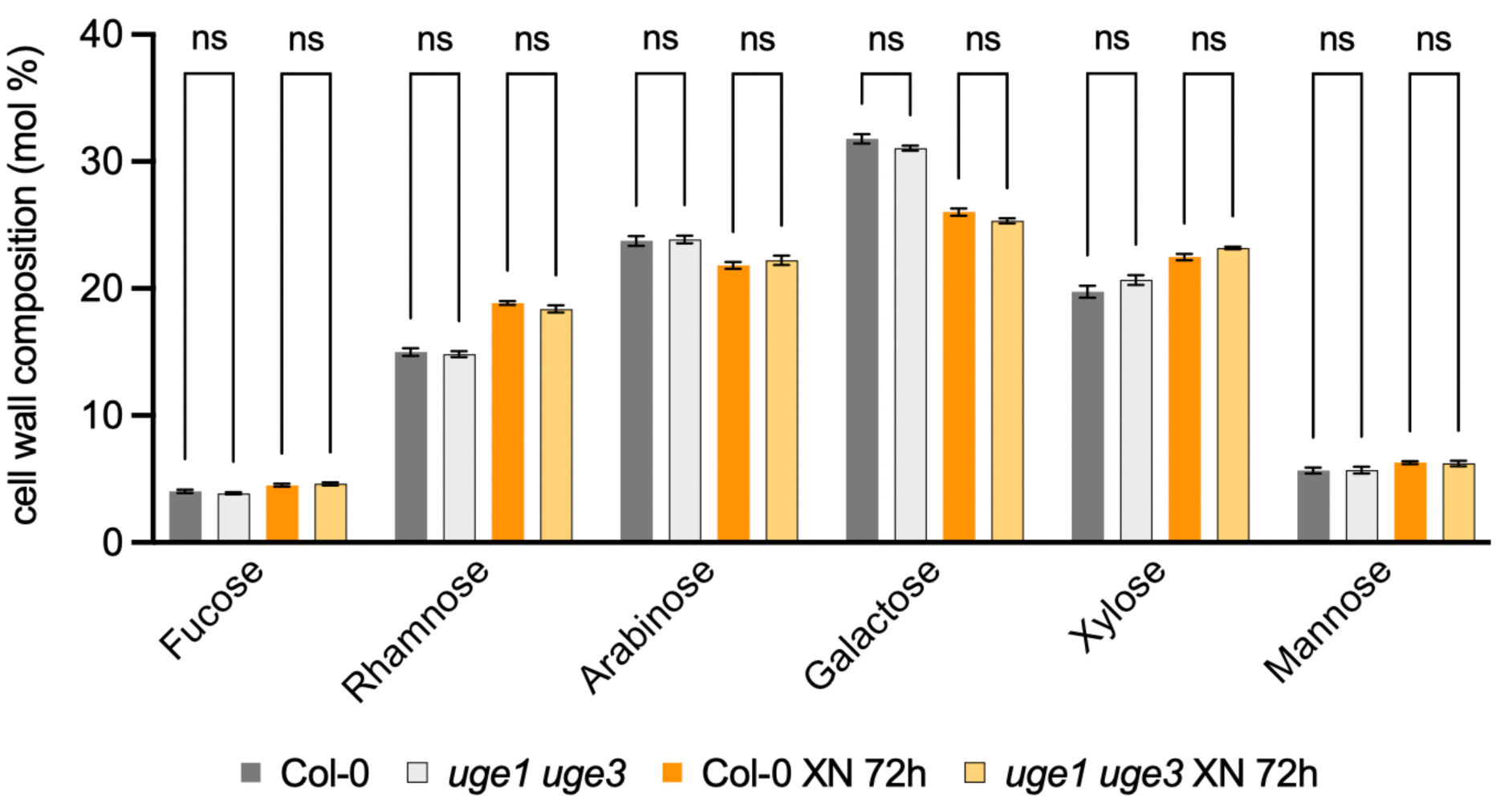
Cell wall monosaccharide composition in *uge1 uge3* rosettes upon starvation. The relative composition of neutral cell wall monosaccharides in whole rosettes of Col-0 and *uge1 uge3* under control conditions (EN) and after 72h XN is depicted as the molar percentage of total cell wall neutral monosaccharide content (mol %). Values are means (n=4) and error bars represent the SEM. A two-way ANOVA and Tuckey’s multiple comparisons test were performed to investigate diXerences between Col-0 and *uge1 uge3* (α = 0.05, ns: not significant).

**Figure S5.**
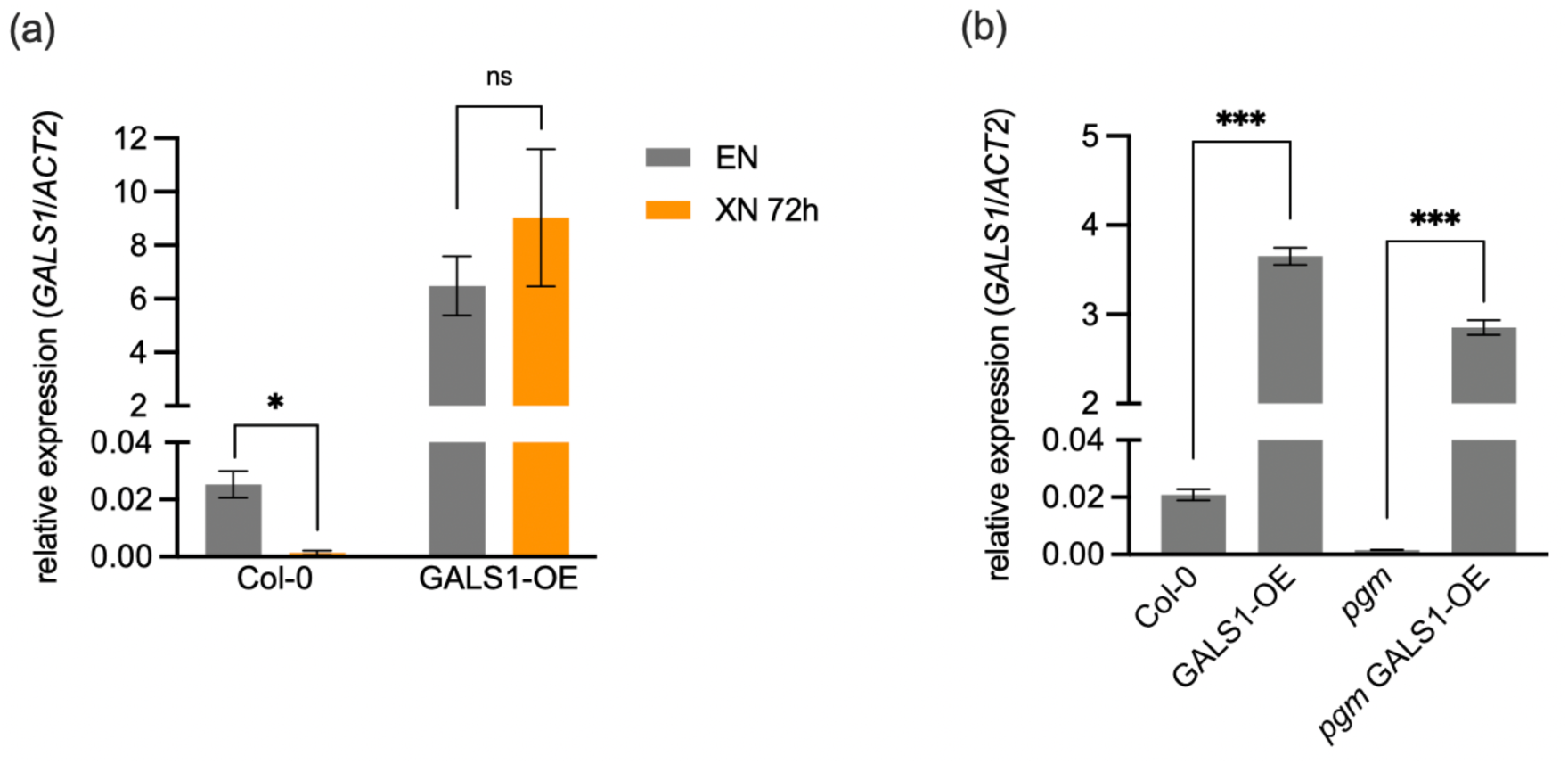
: *GALS1* expression in GALS1-OE lines. (a) The relative expression of *GALS1* was determined in rosette leaves of Col-0 and GALS1-OE at the end of the night (EN) and after 72h extended night (XN). Values are means (n=4) and error bars represent the SEM. Asterisks indicate statistically significant diXerences between XN 24h and EN according to a Student’s t- Test (*p<0.05, ns: not significant). (b) The relative expression of *GALS1* was determined in rosette leaves of Col-0, GALS1-OE, *pgm* and *pgm* GALS1-OE at EN. Values are means (n=3) and error bars represent the SEM. Asterisks indicate statistically significant diXerences between GALS1-OE and their respective genetic background according to a Student’s t-Test (***p<0.001).

**Figure S6.**
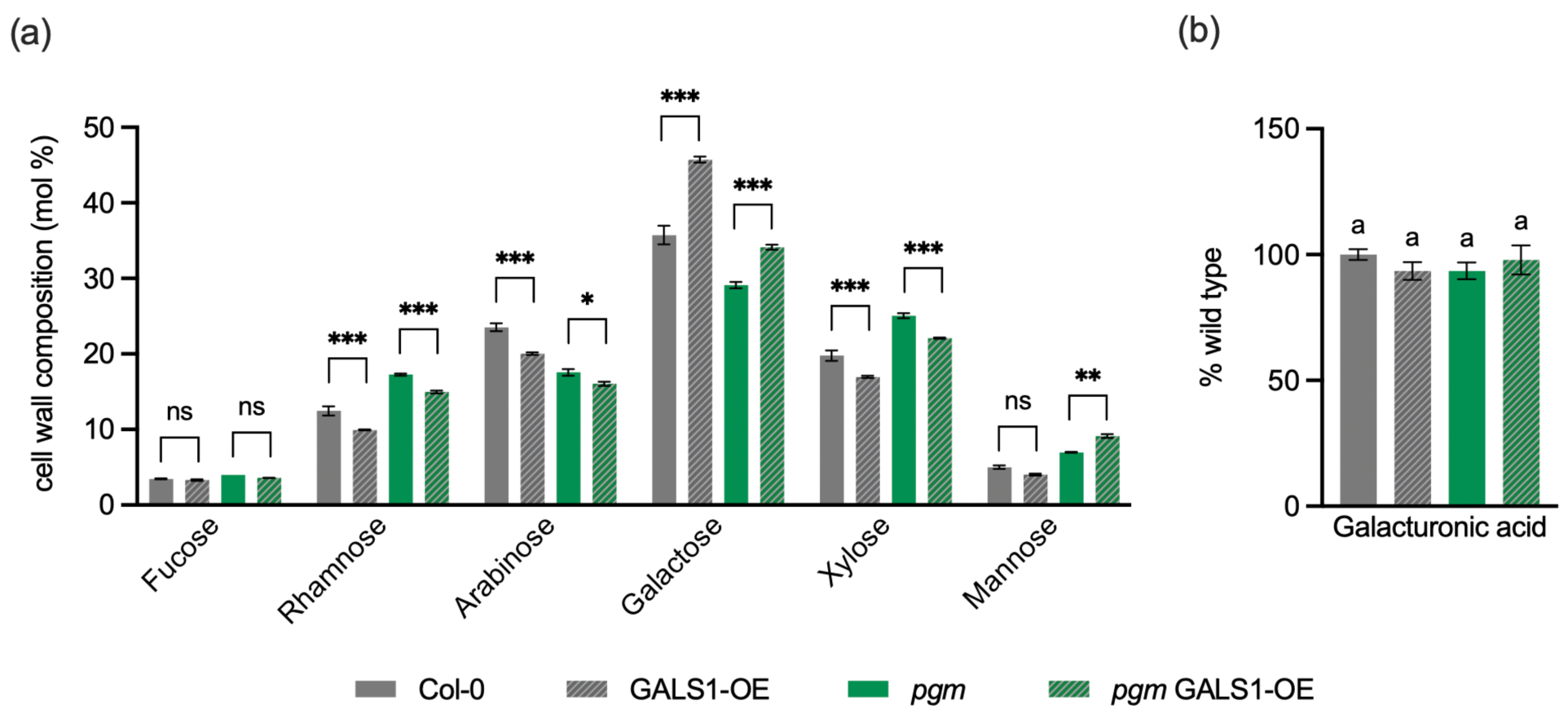
: Cell wall monosaccharide composition in rosettes of GALS1-OE lines. (a) The relative composition of neutral cell wall monosaccharides in whole rosettes of Col-0, GALS1-OE, *pgm* and *pgm* GALS1-OE at the end of a 12h night (EN) is depicted as the molar percentage of total cell wall neutral monosaccharide content (mol %). Values are means (n=4) and error bars represent the SEM. Asterisks indicate statistically significant diXerences between GALS1-OE and their respective genetic background for each monosaccharide according to two-way ANOVA and Tuckey’s multiple comparisons test (*p<0.05, **p<0.01, ***p<0.001, ns: not significant). The data for galactose mol % from this dataset is also shown in Fig. 4d. (b) Galacturonic acid content of Col-0, GALS1-OE, *pgm* and *pgm* GALS1-OE rosettes at EN was quantified relative to Col-0. Values are means from two independent experiments (n=9) and error bars represent the SEM. DiXerent letters indicate statistically significant diXerences according to one-way ANOVA and Tukey’s multiple comparisons test (α = 0.05).

**Table S1.** Expression of *BETA-GALACTOSIDASE* (*BGAL*) genes upon nocturnal and dark-induced starvation. Relative gene expression (fold change) of *BGAL* genes in *pgm* rosette leaves compared to Col-0 4, 8 and 12h after start of the night (N) in a 12h/12h (L/D) cycle and in Col-0 rosette leaves after 2, 4, 6, 8, 24 and 48h of extended night (XN) compared to the end of the regular dark phase. Microarray data were taken from Usadel *et al*. (2008). Fold changes larger than 2 are highlighted in bold.

| Gene | AGI | pgm<br>N 4h | pgm<br>N 8h | pgm<br>N<br>12h | Col-0<br>XN<br>2h | Col-0<br>XN<br>4h | Col-0<br>XN<br>6h | Col-0<br>XN<br>8h | Col-0<br>XN<br>24h | Col-0<br>XN<br>48h |
| --- | --- | --- | --- | --- | --- | --- | --- | --- | --- | --- |
| **BGAL1** | AT3G13750 | **5,89** | **5,24** | **2,62** | **2,06** | **2,47** | **2,63** | **2,32** | **2,01** | 1,67 |
| **BGAL2** | AT3G52840 | **2,74** | **2,14** | 1,50 | 1,56 | **2,31** | **2,35** | **2,13** | 1,69 | 1,04 |
| BGAL3 | AT4G36360 | 0,54 | 0,30 | 0,20 | 0,99 | 0,65 | 0,45 | 0,33 | 0,18 | 0,07 |
| **BGAL4** | AT5G56870 | **13,83** | **24,07** | **21,17** | **5,80** | **22,19** | **37,87** | **31,97** | **22,38** | **24,33** |
| BGAL5 | AT1G45130 | 0,94 | 0,56 | 0,54 | 1,04 | 0,84 | 0,64 | 0,93 | 1,31 | 0,75 |
| **BGAL6** | AT5G63800 | 1,71 | 1,64 | 1,39 | 1,19 | **2,03** | 1,81 | **2,07** | **2,13** | **2,31** |
| BGAL7 | AT5G20710 | 0,93 | 0,86 | 1,08 | 0,93 | 0,94 | 0,94 | 0,93 | 1,14 | 1,11 |
| BGAL8 | AT2G28470 | 0,48 | 0,39 | 0,28 | 1,01 | 1,17 | 1,26 | 1,05 | 0,45 | 0,26 |
| BGAL9 | AT2G32810 | 0,83 | 0,76 | 0,68 | 0,86 | 0,84 | 0,87 | 0,67 | 0,54 | 0,37 |
| **BGAL10** | AT5G63810 | 0,76 | 0,96 | 0,69 | 0,90 | 1,63 | **4,25** | **8,00** | **4,63** | **2,88** |
| BGAL11 | AT4G35010 | 0,91 | 0,90 | 1,03 | 0,94 | 0,93 | 0,85 | 0,97 | 0,98 | 1,07 |
| BGAL12 | AT4G26140 | 1,24 | 1,39 | 1,34 | 1,68 | 1,59 | 1,83 | 1,72 | 1,71 | 1,55 |
| BGAL13 | AT2G16730 | 0,86 | 0,93 | 0,95 | 0,88 | 0,95 | 0,99 | 1,13 | 1,02 | 0,91 |
| BGAL14 | AT4G38590 | 0,87 | 1,01 | 0,97 | 1,21 | 1,00 | 1,03 | 1,04 | 1,07 | 1,16 |
| BGAL16 | AT1G77410 | 0,73 | 0,85 | 1,04 | 0,83 | 0,84 | 0,90 | 0,95 | 1,01 | 1,10 |
| BGAL17 | AT1G72990 | 0,75 | 0,88 | 1,04 | 1,08 | 0,98 | 0,96 | 1,00 | 0,79 | 1,15 |

**Table S2.**
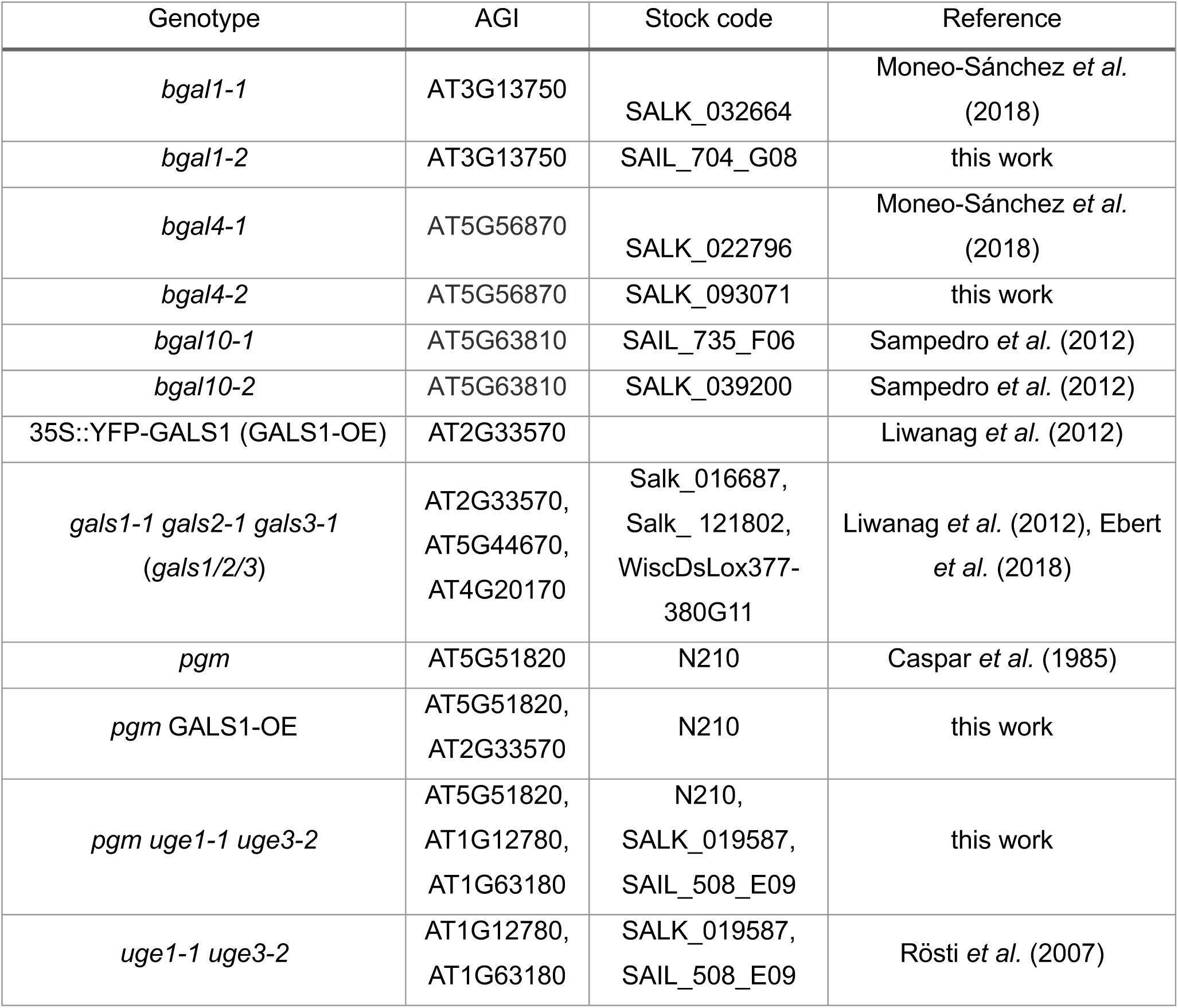
Arabidopsis genotypes used in this study.

| Genotype | AGI | Stock code | Reference |
| --- | --- | --- | --- |
| *bgal1-1* | AT3G13750 | SALK_032664 | Moneo-Sánchez *et al.* (2018) |
| *bgal1-2* | AT3G13750 | SAIL_704_G08 | this work |
| *bgal4-1* | AT5G56870 | SALK_022796 | Moneo-Sánchez *et al.* (2018) |
| *bgal4-2* | AT5G56870 | SALK_093071 | this work |
| *bgal10-1* | AT5G63810 | SAIL_735_F06 | Sampedro *et al.* (2012) |
| *bgal10-2* | AT5G63810 | SALK_039200 | Sampedro *et al.* (2012) |
| 35S::YFP-GALS1 (GALS1-OE) | AT2G33570 |  | Liwanag *et al.* (2012) |
| *gals1-1 gals2-1 gals3-1*<br>(*gals1/2/3*) | AT2G33570,<br>AT5G44670,<br>AT4G20170 | Salk_016687,<br>Salk_121802,<br>WiscDsLox377-380G11 | Liwanag *et al.* (2012), Ebert<br>*et al.* (2018) |
| *pgm* | AT5G51820 | N210 | Caspar *et al.* (1985) |
| *pgm* GALS1-OE | AT5G51820,<br>AT2G33570 | N210 | this work |
| *pgm uge1-1 uge3-2* | AT5G51820,<br>AT1G12780,<br>AT1G63180 | N210,<br>SALK_019587,<br>SAIL_508_E09 | this work |
| *uge1-1 uge3-2* | AT1G12780,<br>AT1G63180 | SALK_019587,<br>SAIL_508_E09 | Rösti *et al.* (2007) |

**Table S3.**
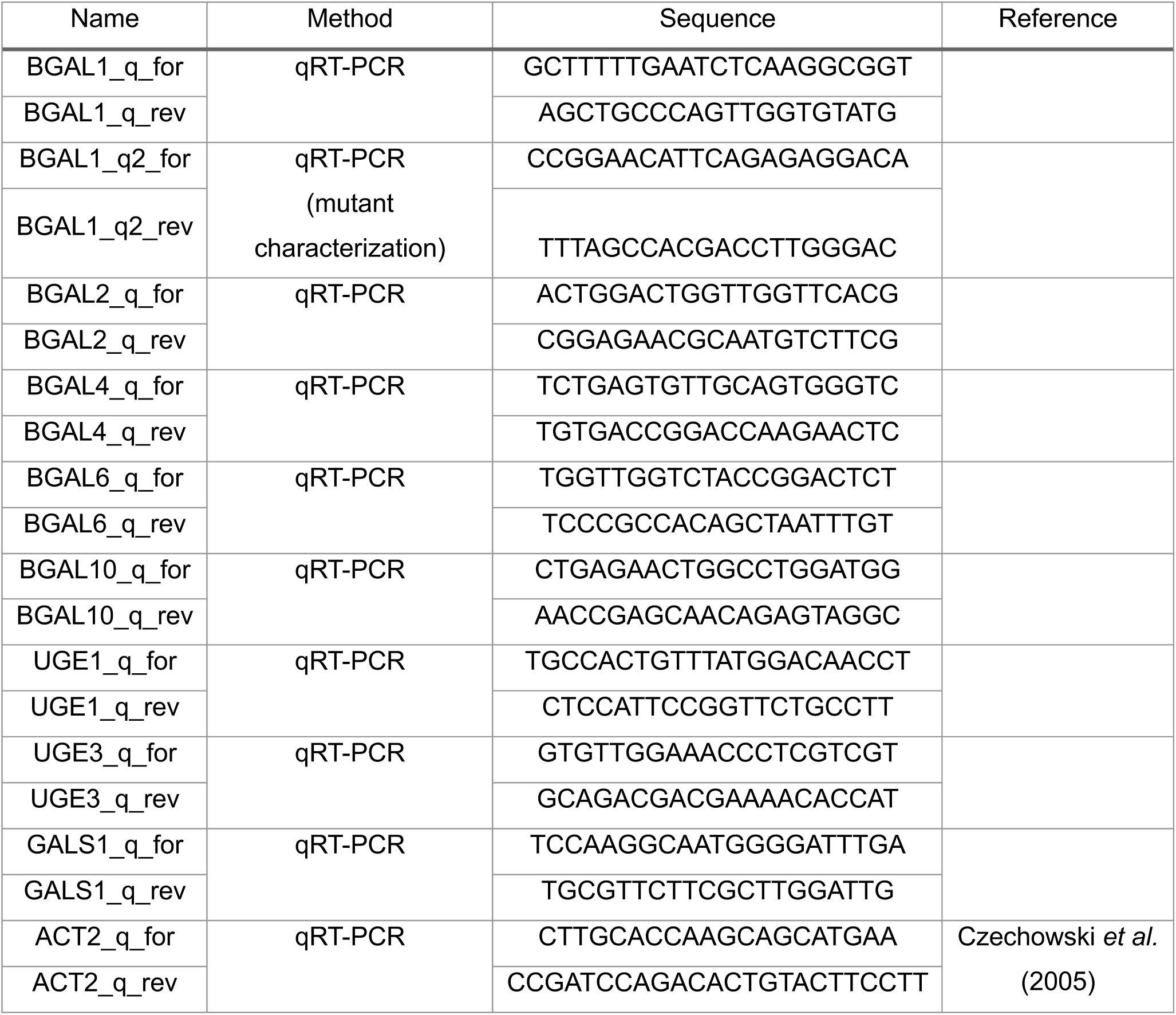
Primers used in this study.

